# *Brucella abortus* histidine auxotrophs are copper sensitive

**DOI:** 10.1101/2025.10.29.685345

**Authors:** Charline Focant, Agnès Roba, Elisabeth Wanlin, Katy Poncin, Xavier De Bolle

**Affiliations:** URBM, Department of Biology, Narilis, University of Namur, Namur, Belgium

**Author notes:** Corresponding author 61 rue de Bruxelles, 5000 Namur, Belgium, Website : www.urbm.be.

## Abstract

Despite decades of investigation into bacterial pathogens, the conditions met by intracellular bacteria are still unclear. These conditions can include access to nutrients, such as amino acids, and exposure to toxic compounds, like copper. To investigate the ability of *Brucella abortus*, a facultative intracellular pathogen responsible for a major zoonosis, to cope with copper, we performed a Tn-seq analysis to identify copper-sensitive mutants. Unexpectedly, we realized that classical copper resistance systems (involving CopA and CueO homologs) do not appear to be robustly needed, while histidine and purine biosynthesis pathways are crucial to cope with copper. We show that *hisA, hisB, hisC* and *hisD* mutants are auxotrophic for histidine and sensitive to copper. This suggests that the reported attenuation of *his* mutants in macrophages could be based on auxotrophy or/and copper sensitivity. Therefore, we generated suppressor strains with a restored resistance to copper for *hisC*, but still auxotrophs for histidine. Our data suggest that this suppression is due to the overproduction of a homolog of OppA, a periplasmic oligopeptide binding protein. Analysis of these suppressors shows that the absence of histidine biosynthesis capacity, and not copper sensitivity, is required for optimal growth of *B. abortus* in macrophages.

## Introduction

*Brucella* spp. are Gram-negative bacteria that belong to the Hyphomicrobiales order, which is part of the alpha-proteobacteria class (1,2). Bacteria from the genus *Brucella* are intracellular, facultatively extracellular pathogens responsible for brucellosis (3). This worldwide zoonosis disease affects a wide range of hosts, including domestic animals and humans, illustrating the complexity of *Brucella* spp. (4). The tropism of the bacteria for the reproductive systems correlates with abortion and sterility as main symptoms in animals while humans present undulant-fever symptoms (3,5,6). *Brucella* employs sophisticated strategies to enter, survive and proliferate within host cells - encompassing both professional phagocytes and non-phagocytic cells - thereby effectively ensuring its pathogenicity (3,6). Once within the cell, *Brucella* is found enclosed in a membrane-bound compartment, the *Brucella*-containing vacuole (BCV), which traffics along the endocytic pathway and undergoes an acidification of the compartment, becoming the endocytic BCV (eBCV) (7,8). The vacuole acidification is crucial to produce the *Brucella* VirB Type IV secretion system (7,9), which secretes effector proteins inside the cell (10,11). The establishment of the replicative niche (rBCV) inside the endoplasmic reticulum (ER) is dependent on the VirB function (7,8). As a last step, the rBCVs are converted into autophagic BCVs (aBCVs) by getting associated to atypical autophagic membranes (12). Inside the aBCVs, *Brucella* can egress from the cell, being therefore able to reinfect the surrounding cells (12). During its trafficking, *Brucella* probably encounters many stresses, most of which are unknown or indirectly deduced from attenuated mutants.

In the arms race of host-pathogen co-evolution, host cells use several strategies, one of which is the sequestration of key metal ions for the pathogenic bacteria (13). Iron deprivation by host cells is well established. Most iron ions are predominantly sequestered, e. g. bound by transferrin and hemoglobin, rendering them unavailable to bacteria (14,15). Neutrophils recruited to the infection site release calprotectin which sequesters manganese and zinc (16,17). Manganese and iron can also be depleted from the phagosomes through the natural resistance-associated macrophage protein 1 (NRAMP1) in macrophages and neutrophils (13,18). Despite the well-established role of metal starvation as a defence mechanism against pathogens, it is becoming increasingly evident that metal intoxication also occurs during infection to fight invading pathogen (16,19). Host cells exploit copper toxicity to poison intrusive pathogens (20). In response to interferon-γ (IFN-γ) stimulation, activated macrophages accumulate copper within phagolysosomes (21). IFN-γ induces the expression of the copper transporter CTR1, allowing copper import into the cytosol (21,22), where it is delivered by the ATOX1 chaperone to the ATP7A transporter associated to the phagolysosomal membrane (21,23–25) able to pump copper into the lumen of the phagosome. ATP7A expression is also upregulated by IFN-γ (24). Copper resistance has been linked to the virulence of intravacuolar pathogens such as *Mycobacterium tuberculosis* (25,26) and *Salmonella enterica* (27).

Several defensive mechanisms are used by bacteria to combat toxic concentrations of metals. A simple mechanism of averting metal toxicity is to sequester metal ions within the periplasm or the cytoplasm, resulting in fewer than one copper atom being available per cell (15,28,29). When metal ions accumulate to toxic levels, bacteria induce the expression of specific adapted genes. This defence response includes the induction of eflux system as well as proteins involved in metal sequestration and storage (16). The bacterial response to copper could involve a core *cue* regulon, which is relatively well conserved among proteobacteria (28). The *cue* regulon includes CueR, a Copper-responsive metalloregulatory protein which up-regulates *cueO* and *copA* expression upon increased intracellular copper concentration (30). CueO, the periplasmic multicopper oxidase, oxidizes cuprous ion to cupric ion, which is less harmful (31). CopA is a Copper efflux P-type ATPase (32). A cuprochaperone, CopZ, is also part of the Cue eflux system (33). Most of *S. enterica* serotype possess a duplication of the cue regulon, known as the gol regulon (34,35). The expanded anti-copper arsenal of *S. enterica* is presumably an adaptation for survival within macrophages (28,34). A second copper eflux system, the Cus system, is also present in *Escherichia coli* and *S. enterica*. This second regulon is controlled by the two-component system CusRS (28), which activates the expression of the *cusCFBA* operon in response to elevated copper levels (31). The *cusCBA* genes encode the CusCBA complex, a resistance, nodulation and division (RND) proton-cation antiporter (28,31), while *cusF* encodes a periplasmic cuprochaperone (28).

The specific environmental conditions encountered by intracellular bacteria remain poorly characterized. These conditions may involve limited access to nutrients, such as amino acids, and exposure to compounds like copper. To explore how *B. abortus* responds to copper stress, we conducted a transposon sequencing (Tn-seq) analysis to identify mutants with increased copper sensitivity. Although the classical CueO and CopA homologs do not appear to be required for *B. abortus* resistance to copper, the histidine biosynthesis pathway was found to be necessary for growth in the presence of copper. Since it is known (and confirmed here) that *his* mutants are attenuated in macrophages, this phenotype could be due to auxotrophy for histidine, and/or to sensitivity to copper. By generating suppressors with a restored copper resistance, we show that the attenuation of the *his* mutants in macrophages is probably not related to copper sensitivity.

## Results

### Identification of genes involved in copper resistance in *B. abortus*

The mechanism by which *Brucella* copes with copper toxicity remains unknown. To identify genes required to face copper stress, a Tn-seq was performed on *B. abortus* 544 using a mini-Tn*5* (36). To select the appropriate copper concentration for the Tn-seq, *B. abortus* and *E. coli* mini-Tn*5* carrier strains were first mated. After incubation and OD normalization, a serial 10-fold dilutions were spotted on TSB agar plates supplemented with an increasing CuSO_4_ concentrations CFUs were then counted to identify the optimal copper concentration. A final concentration range of 1.6 to 2 mM CuSO_4_ was chosen for the selection corresponding to the concentration where WT was not impacted. Once the copper concentration to use for the Tn-seq was determined, the mating was repeated, and the *Brucella* library was grown on rich medium for the control condition, and 2 mM CuSO_4_ supplemented media for the test condition. From the library, 4.8 x 10^6^ random mutants were recovered and sequenced using a deep sequencing method. We identified 1 684 839 and 1 743 143 unique insertion sites for control and copper condition respectively, illustrating genome saturation with a unique insertion every 1.95 bp for control and every 1.89 bp for the stress condition, on average. The raw data were analysed by an automatic process called TnBox, following a previously described method (37). The transposon insertion frequency (TnIF) for each gene was computed according to the following definition: TnIF is defined as the per-gene average of log_10_(*r*+1)/*l*, where *r* is the number of miniTn*5* insertions at a given nucleotide and *l* is the coding sequence length. The calculation was performed considering only the central 80% of the coding sequence (37). A TnIF value below the genome-wide average indicates that disruption of the gene decreases bacterial fitness. Then, the ΔTnIF was determined as the difference between the TnIF under copper stress condition and the TnIF under control conditions (ΔTnIF = TnIF*copper* - TnIF*control)*. A gene will be annotated as required to grow in the presence of copper if the ΔTnIF is lower than 2 times the standard deviation of all ΔTnIF values (ΔTnIF > - 0,357). Through the Tn-seq analysis, thirty-eight genes were identified as essential for growth under copper-supplemented condition (Table 1).

**Table 1.**
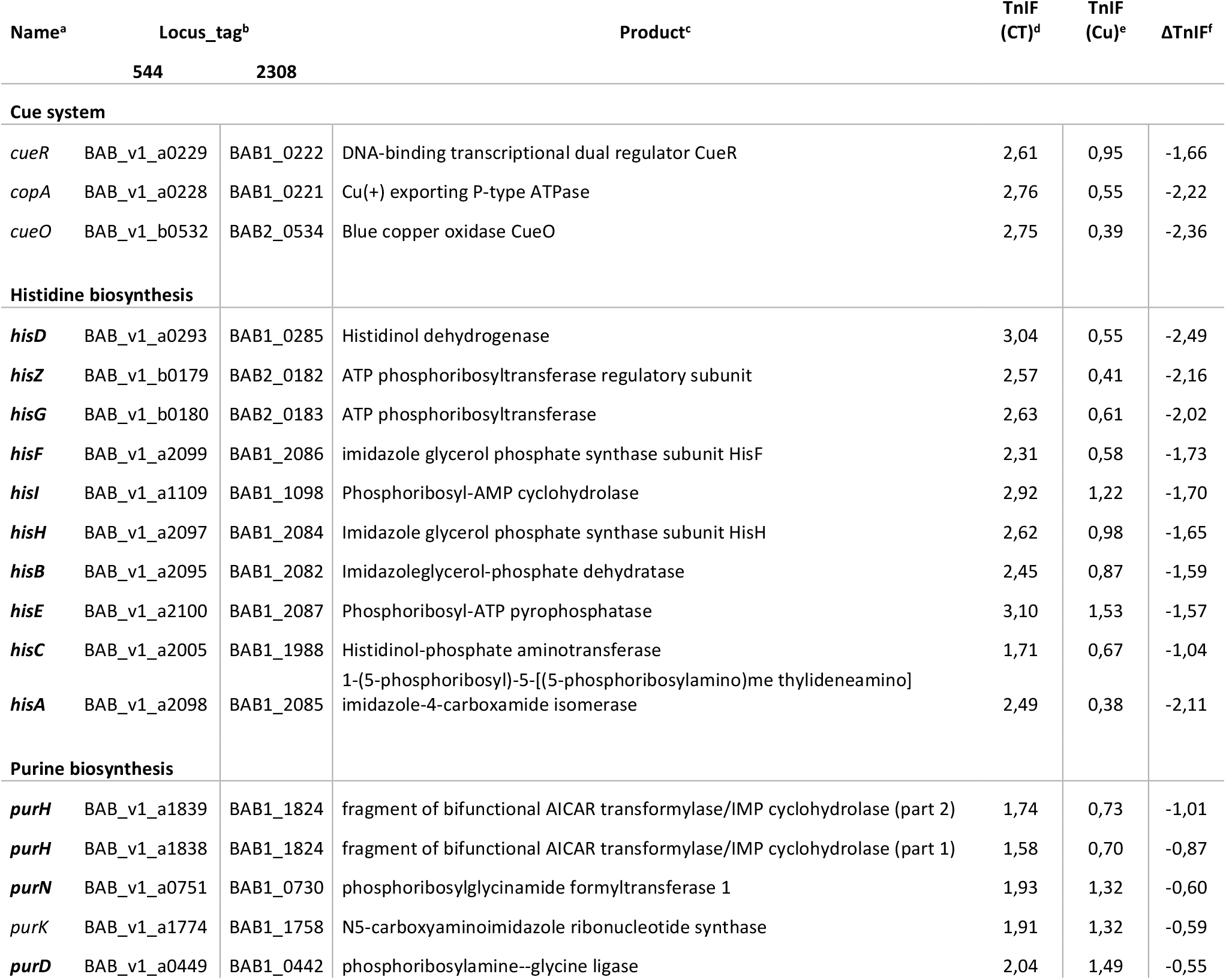
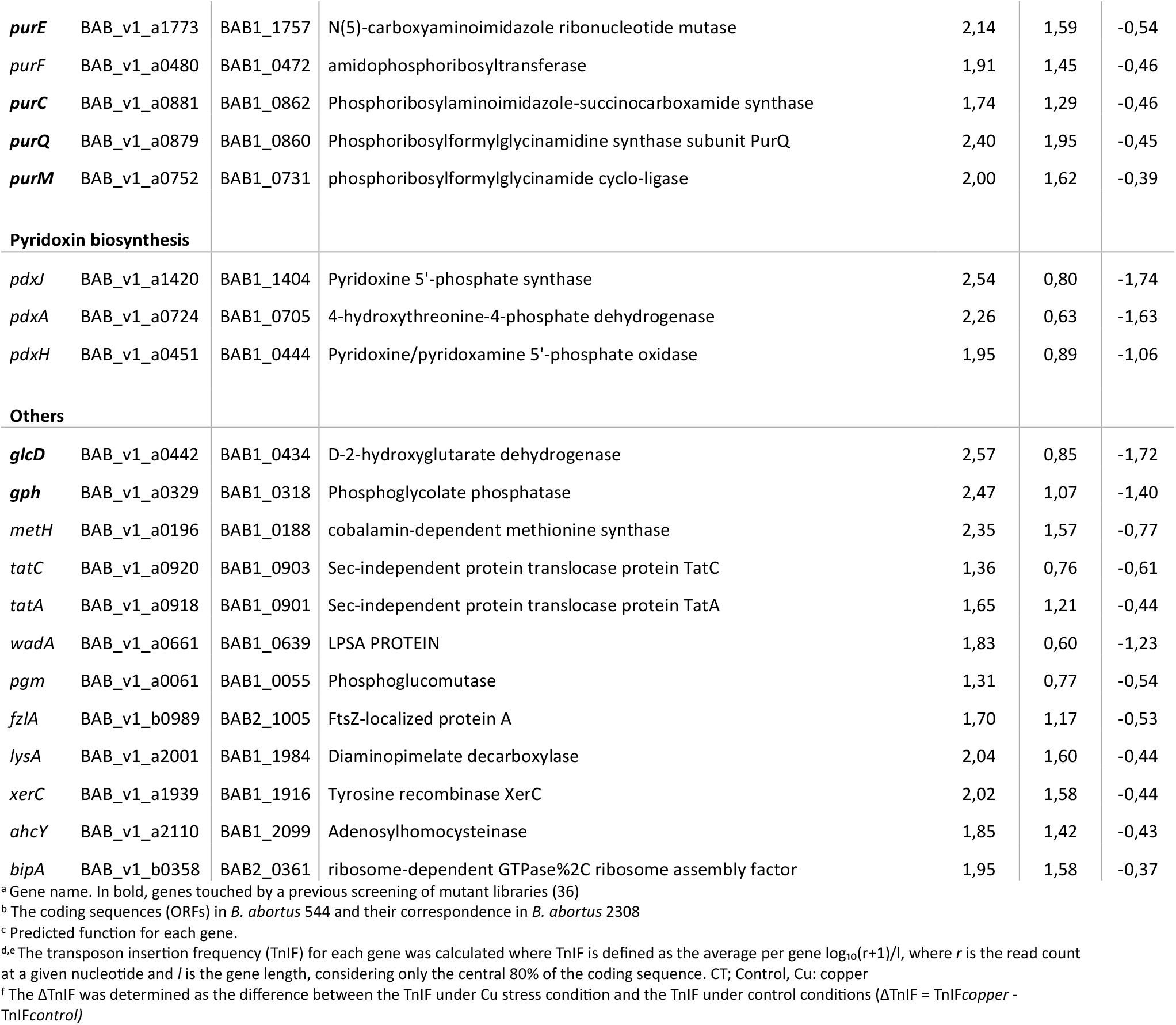
Attenuated *B. abortus* mutants on copper 2 mM-supplemented TSB plates.

The genes coding for the Cue system, *copA, cueO* and the regulator *cueR*, involved in copper resistance, appeared to be essential in the Tn-seq copper condition (Table 1), as expected compared to the data available from the work in *E. coli* (31). CopA is an efflux P-type ATPase of the *cue* regulon while CueO is a periplasmic multi copper oxidase (32). The expression of both *copA* and *cueO* is regulated by the copper-responsive transcriptional regulator CueR (30). The *cue* regulon also includes *copZ* (BAB1_0960 in *B. abortus* 2308 and BAB_v1_a0976 in *B. abortus* 544 (33)) which is not scored as required in the Tn-seq copper condition.

Genes coding for the whole histidine biosynthesis pathway (*hisA, hisB, hisC, hisD, hisE, hisF, hisG, hisH* and *hisI*) are highlighted in our Tn-seq to be required to cope with copper stress. Each step of the pathway is catalysed by a specific enzyme where the five first steps lead to the production of imidazole glycerol phosphate (IGP) and 5-aminoimidazole-4-carboxamide ribonucleotide (AICAR), an intermediate in purine biosynthesis (See Fig. S1). Histidine is produced from IGP after four other enzymatic reactions (See Fig. STba1). While in *E. coli* and *S. enterica* Typhimurium, the his pathway is genetically organised in a single operon, it is distributed across both chromosomes in *Brucella* spp. (38).

Linked to the histidine biosynthesis pathway, the purine *de novo* synthesis pathway appeared to be also crucial to face copper stress. Indeed, most of the genes - *purH* (part I and II), *purN, purK, purD, purE, purF, purC, purQ* and *purM* - required for inosine monophosphate (IMP) production were found to be required in the presence of copper.

Next, three genes from the vitamin B6 biosynthesis (*pdxJ, pdxA* and *pdxH*) also emerged from our analysis as being required. These genes code for enzymes allowing the biosynthesis of vitamin B6, in its active form pyridoxal 5’-phosphate (39). Vitamin B6 is utilized as a cofactor by the HisC enzyme (40), which may explain why it is required to face copper stress.

Finally, there were twelve other genes that also presented a ΔTnIF lower than 2 times the standard deviation. Among them, *glcD, tatC, pgm* and *fzlA* were already essential in the control condition, while *metH, tatA, wadA, lysA, xerC, ahcY* and *bipA* had low fitness in the control condition. Only *gph* did not present any impact on growth in the absence of copper.

### CopA and CueO are not required for copper resistance in *Brucella abortus*

To date, the mechanisms underlying copper resistance in *B. abortus* remain largely uncharacterized. Only few homologs of genes involved in copper homeostasis are predicted from the *Brucella* genomes, including *copA, cueO* and *cueR* (41). These three genes were identified as required under copper stress condition in our Tn-seq screen (Table 1). Given their predicted function in copper transport and detoxification, we first focused on testing the role of *copA* and *cueO* in *B. abortus* copper resistance by constructing markerless single-gene deletion mutants called Δ*copA* and Δ*cueO*. The growth of Δ*copA* and Δ*cueO* was monitored during 48 hours in liquid culture in rich medium in the presence of 2 mM CuSO_4_ (Fig. 1B). Unexpectedly, both mutants did not present any sensitivity to copper stress, displaying growth curves similar to the wild-type (WT) strain. Δ*copA* presented a slight growth defect in the control condition (Fig. 1A) compared to the WT which was also found in the copper condition (Fig. 1B). The minor effect of copper on Δ*cueO* was not confirmed when the copper concentration was increased.

**Figure 1.**
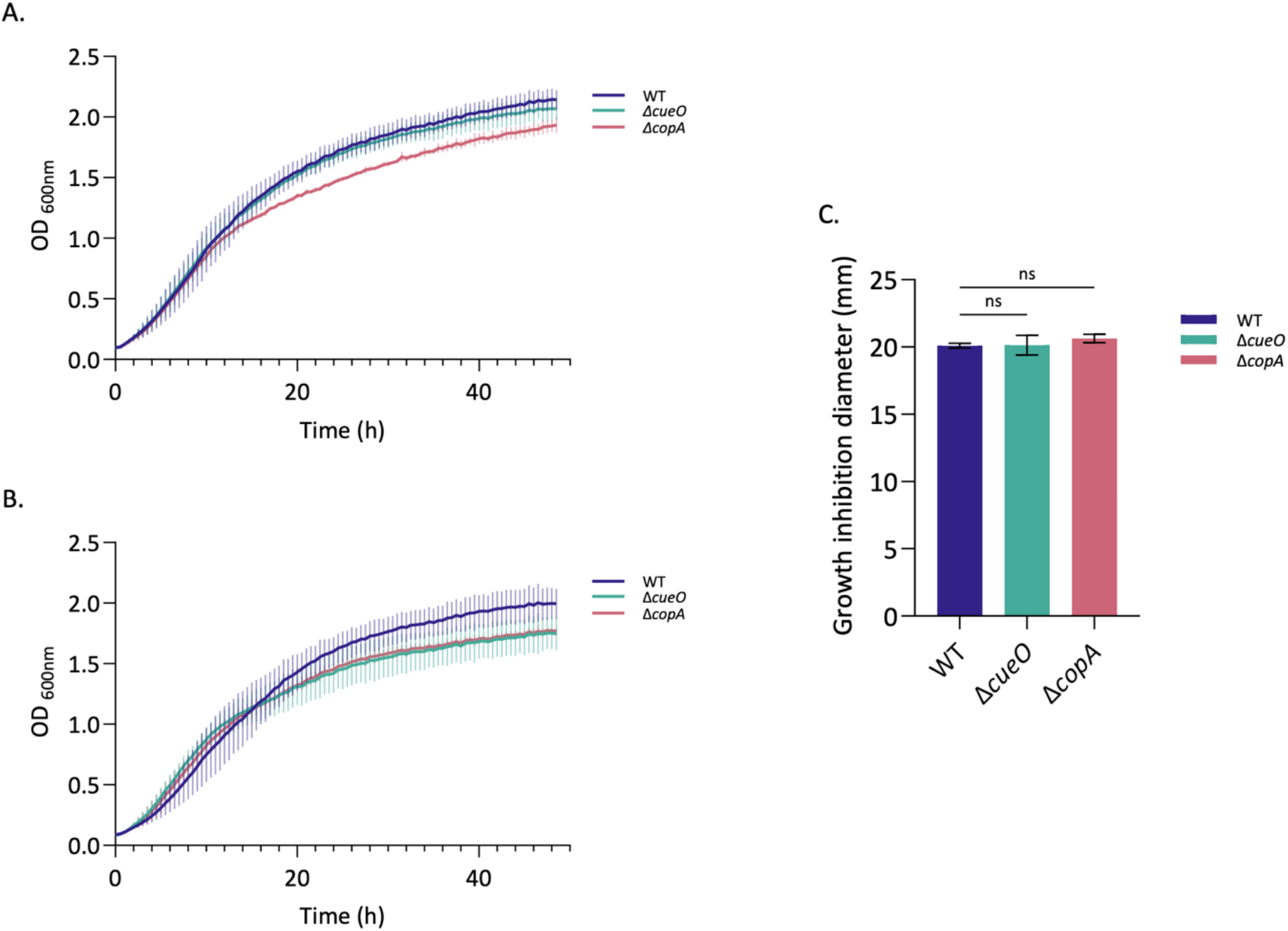
The *copA* and *cueO* genes are not required for copper resistance. WT, Δ*copA* and Δ*cueO* strains were tested for their sensitivity to copper toxicity for growth. **A**. WT, Δ*copA* and Δ*cueO* strains were grown in liquid TSB rich medium. **B**. WT, Δ*copA* and Δ*cueO* strains were grown in TSB rich medium containing 2 mM CuSO_4_. **A**., **B**. The optical density (OD) at 600 nm was measured every 30 minutes for 48 hours. Data represent three independent replicates. **C**. Five hundred μl of an overnight culture of each *B. abortus* strain normalised to OD 0.1 were added to 4.5 ml of TSB soft agar (0.7% agar) and then plated on a TSB plate. In the middle, a well was dug and filled with 100 μl of 200 mM CuSO_4_. Inhibition zones around the agar well were measured after 3 days of incubation at 37°C and the average diameter (± standard deviation, SD) reported in a histogram. The data represent the mean ± SD and were compiled from three independent replicates. Statistical analysis was carried out by an ANOVA one-way followed by Dunnett’s multiple comparisons test (n.s., non-significant, *P* > 0.05).

We also conducted a soft agar well diffusion assay where the well was filled with 100 μl of 200 mM CuSO_4_. The inhibition zone was measured around the well and reported in millimetres, with the results presented as a bar graph. As with the growth curves, Δ*copA* and Δ*cueO* displayed the same inhibition zone as the WT strain around the copper source, confirming the absence of sensitivity of the two mutants in these conditions (Fig. 1C). Taken together, these results demonstrate that the deletion of *copA* or *cueO* does not impact copper resistance in *B. abortus*, contrary to what has been shown in *E. coli*, and also in sharp contrast to our Tn-seq data (Table 1).

### The *his* mutants are auxotrophic for histidine and sensitive to copper

In addition to our Tn-seq results, histidine biosynthesis genes have been already unveiled as required in *B. abortus* during RAW 264.7 macrophages infection (36), as well as previous studies with *B. suis* (42,43) and Tn-seq with *B. melitensis* in mice (37). A series of deletion mutants in the histidine biosynthesis pathway has been constructed; *hisA, hisB* (previously reported (44)), *hisC* and *hisD*. The auxotroph phenotypes of these four mutants have been confirmed by their inability to proliferate in minimal medium in the absence of amino acids. Their growth was rescued either when the gene was introduced in *trans* on a single-copy plasmid (Fig. 2) or if histidine was provided in the culture medium.

**Figure 2.**
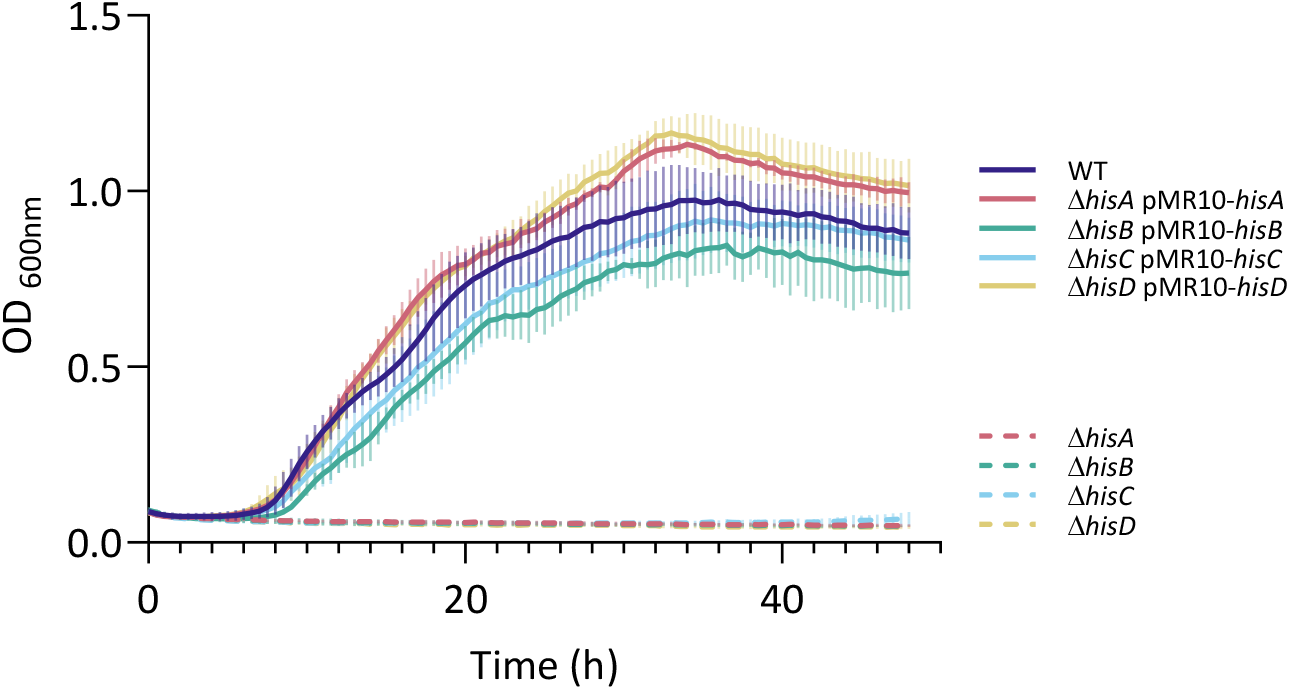
*B. abortus* Δ*hisA*, Δ*hisB*, Δ*hisC* and Δ*hisD* are auxotrophic for histidine. WT and *his* mutant strains were tested for their histidine auxotrophy. Strains were grown in liquid PE minimal medium. OD _600nm_ was measured every 30 minutes for 48 hours. The pMR10 is a low copy plasmid. The data represent three independent replicates.

Since the classical copper homeostasis actors (CopA, CueO) were not required for growth of *B. abortus* in the presence of a copper stress, we decided to investigate the histidine biosynthesis pathway in the context of copper stress. The growth of the *his* mutants in the presence of copper was studied through diverse approaches. First, all strains were cultivated in liquid rich medium with a high copper concentration (1.6 mM) and the optical density (OD) was measured over a period of 48 hours (Fig. 3A), as done with Δ*copA* and Δ*cueO* strains. As suggested by the Tn-seq results, all auxotrophic mutants struggled to grow in the presence of the metal compared to the WT strain, which did not demonstrate a growth defect. The *hisA* mutant was particularly affected. However it already struggled in control condition compared to the other strains (See Fig. S2). The same sensitivity phenotype for all the *his* mutants was observed with 2mM CuCl_2_, indicating that the growth defect was due to copper rather than sulfate. The copper sensitivity of the *his* auxotroph mutants was rescued upon genetic complementation (Fig. 3B). Similar observations were made with the same strains cultivated with a concentration of 2 mM of CuSO_4,_ but with an incomplete complementation (See Fig. S3). These preliminary results suggested that impairment of the histidine biosynthesis pathway leads to a copper sensitivity. The sensitivity of *his* mutants to copper was confirmed by a soft agar well diffusion assay in which the inhibition zone around the copper-containing well was measured. As with growth curves, the copper sensitivity was rescued when the strains were complemented with the missing genes (Fig. 3C). The growth of the four *his* auxotroph mutants was tested on solid TSB medium supplemented with 1.6 or 2 mM CuSO_4_. Results showed a growth defect for each *his* mutant, at the opposite of the WT strain which grew normally (Fig. 4). On these plates, a few colonies among the mutants were observed. The presence of mutations in these colonies could potentially enable them to withstand the presence of copper. In this case, these colonies could be suppressor clones, *i*.*e*. clones in which a second mutation cancels out the phenotypic effect of the first mutation (Δ*hisA*, Δ*hisB*, Δ*hisC* or Δ*hisD*). Identifying these secondary mutations could therefore provide valuable insight to understand how these strains are able to cope with copper.

**Figure 3.**
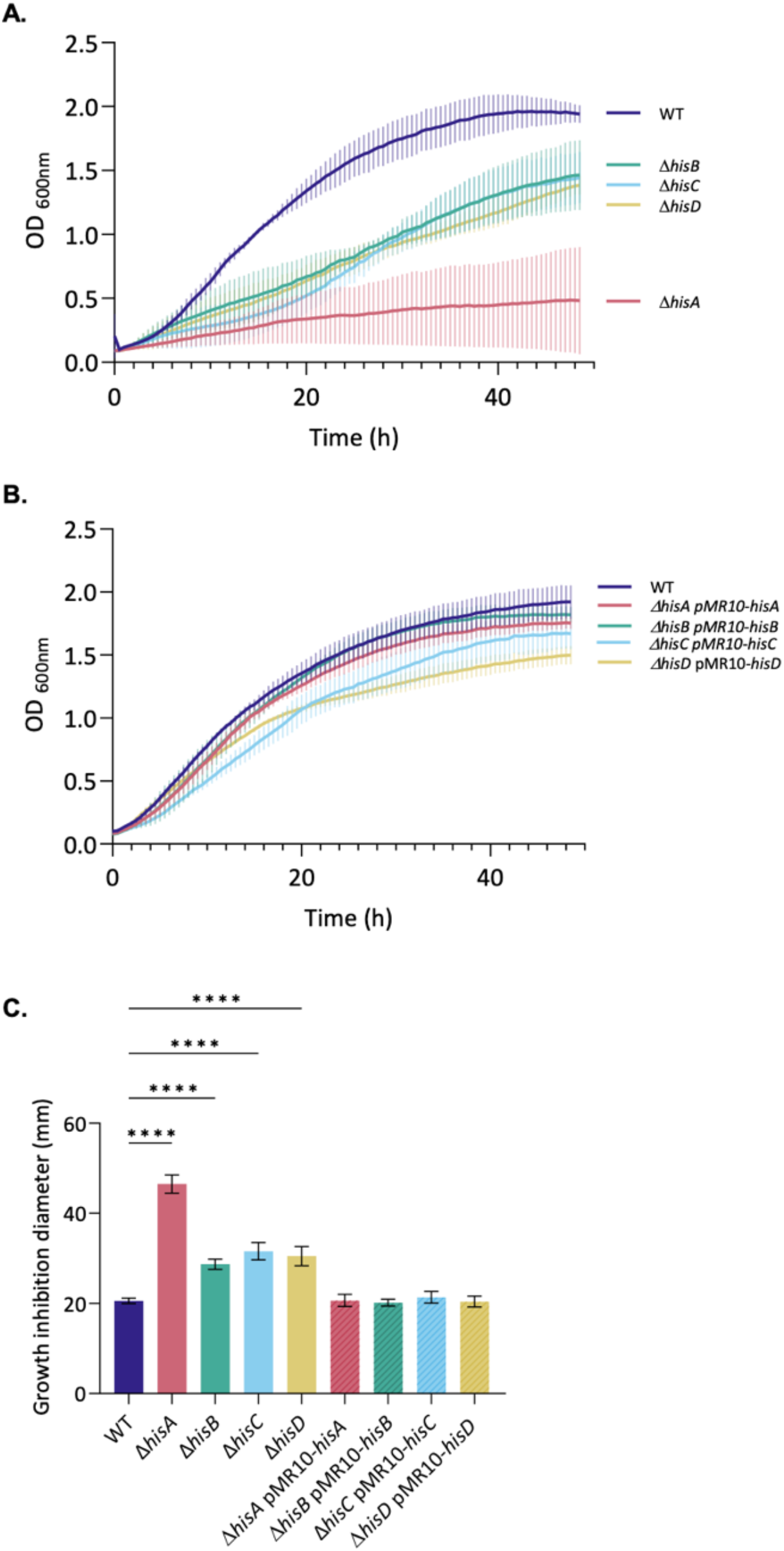
Histidine auxotrophic mutants are copper sensitive. WT, Δ*hisA*, Δ*hisB*, Δ*hisC*, Δ*hisD*, Δ*hisA* pMR10-*hisA*, Δ*hisB* pMR10-*hisB*, Δ*hisC* pMR10-*hisC* and Δ*hisD* pMR10-*hisD* strains were tested for their sensitivity to copper toxicity for growth. **A**. WT, Δ*his* and **B**. complemented strains were grown in liquid TSB rich medium containing 1.6 mM of CuSO_4_. OD _600nm_ were measured every 30 minutes for 48 hours. The data represent three independent replicates. **C**. Experiment performed as in Fig. 1C. Data represent the mean ± SD and were compiled from three independent replicates. Statistical analysis was carried out by an ANOVA one-way followed by Dunnett’s multiple comparisons test (****, *P* <0.0001).

**Figure 4.**
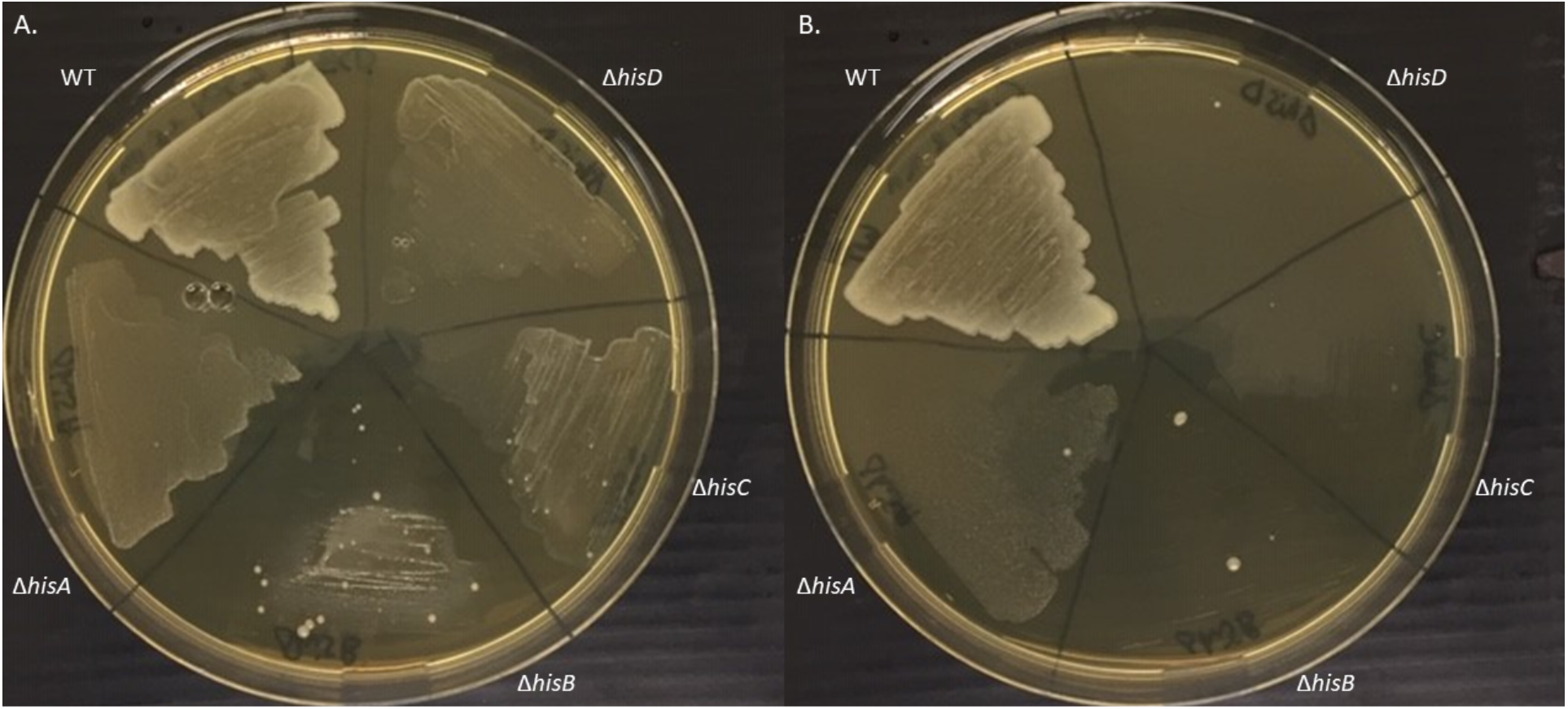
*B. abortus* WT and mutant strains are copper sensitive on plates. WT and Δ*hisA*, Δ*hisB*, Δ*hisC* and Δ*hisD* mutant strains (in this counterclockwise order) was tested for their copper sensitivity on TSB medium containing 1.6 mM (**A**.) or 2 mM (**B**.) of CuSO_4_. Both plates were divided in five parts where 20 μl of each culture were deposited and spread after their OD _600nm_ was normalized to 0.1. Pictures were taken after a four-day incubation at 37°C. The data represent three independent replicates. It is interesting to note the presence of individual colonies for Δ*hisA*, Δ*hisB*, Δ*hisC* and Δ*hisD*, suggesting that suppressors mutants could be obtained for this phenotype.

### Most suppressor mutations affect a pair of periplasmic proteins

In order to verify the suppressor phenotype of the colonies isolated from the copper-supplemented TSB plates, it was necessary to streak each of them onto a new plate with and without copper. Of the 29 potential suppressors isolated, all mutant backgrounds combined, eight were unable to retain the ability to grow on copper-supplemented media. The remaining potential suppressors were able to grow at 2 mM CuSO_4_. The genomic DNA of the remaining 21 copper-resistant clones was extracted and analysed by whole-genome sequencing to identify the mutations that could confer copper resistance in these suppressors. As a control, the genomes of the WT and the parental Δ*hisA*, Δ*hisB*, Δ*hisC* and Δ*hisD* strains were also sequenced. To investigate mutations that were strictly present in the potential suppressor, only the mutations present in the suppressor and not in the parental strain were retained. Strikingly, of the 22 unique mutations identified, 15 appeared within the same operon, the *opp* operon (Table 2). Opp is homologous to an oligopeptide transport system, it is encoded by five genes. The first two genes, *oppA1* and *oppA2*, encode periplasmic proteins. The next two genes, *oppB* and *oppC*, encode permeases, while the final gene, *oppD/F*, encodes an ATP-binding protein. Interestingly, only the two first genes were touched by a mutation in the suppressors (Table 2).

**Table 2.**
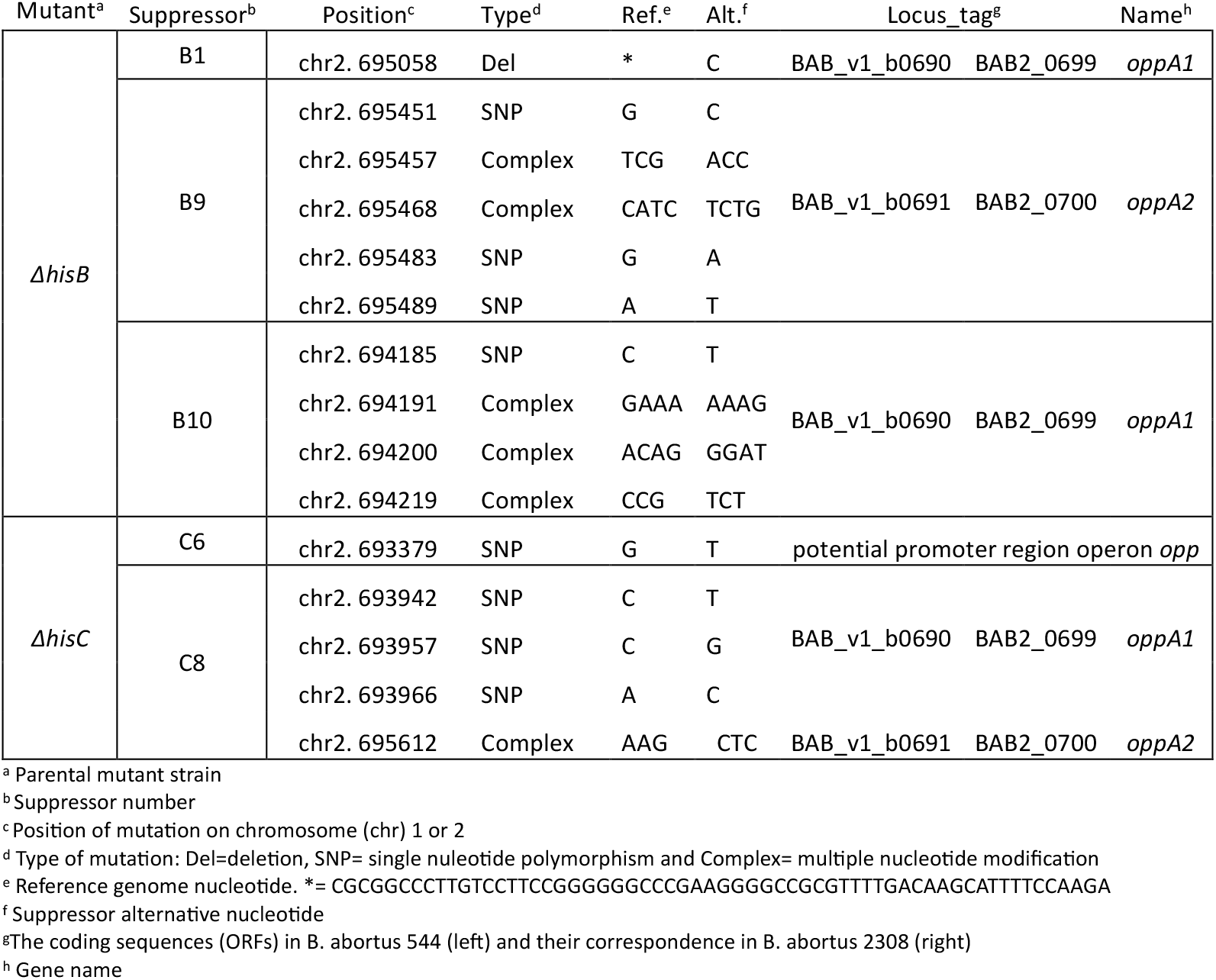
Suppressor mutations.

### The suppressive mutation helps to face copper stress *in vitro*

In this study, we opted to exclude suppressors derived from the *hisB* mutant. Indeed, *hisB* is located downstream of the intersection between the histidine synthesis pathway and purine metabolism, *via* the production of AICAR (Fig. 1). Moreover, in the laboratory, *hisB* mutant had been studied and demonstrated to exhibit a morphological abnormality by the formation of chains, which is associated with a deficiency in cell division (44). It is also noteworthy that suppressors exhibiting a unique mutation within *hisA* or *hisD* background did not affect the *opp* operon (See Table S2). Out of all mutations in the *hisC* mutant background, one stood out as particularly interesting. It is located in the suppressor named C6 (the sixth suppressor isolated within the *hisC* mutant), 70 pb upstream the *oppA1* ATG, in a potential distal promoter region (Table 2). The mutation was investigated as it could influence the expression of the first two coding sequences of the operon.

For the purpose of confirming that the suppressor phenotype is attributable to the identified mutation, it was introduced back into the *hisC* mutant. The potential distal promoter region where the SNP occurred was amplified from C6 genomic DNA and subsequently inserted into Δ*hisC* strain by allelic exchange. The new strain including the C6 mutation was named Δ*hisC*SupPA1 (standing for Suppressor Promoter OppA1). To investigate the hypothesis that the suppressive mutation in the potential promoter enhances the abundance of OppA1 and OppA2 substrate proteins, an overexpression vector for each of the genes was constructed in the pBBR-MCS2 medium-copy plasmid, and introduced into the Δ*hisC* mutant, generating Δ*hisC* pBBR-*oppA1* and Δ*hisC* pBBR-*oppA2* strains, respectively. The histidine auxotroph phenotype was first evaluated by measuring the OD in a minimal media, as previously performed with the *his* mutants. All strains retained the auxotrophic phenotype of the mutant parental strain. Indeed, the growth was rescued when the minimal medium was supplemented with 1 mM of histidine (See Fig. S4).

The suppression phenotype was then studied to confirm its role in the rescue process in the presence of copper stress. For this purpose, the same experiments as for histidine auxotrophic mutants were carried out. The growth of the strains WT, Δ*hisC*, C6, Δ*hisC*SupPA1, Δ*hisC* pBBR-*oppA1* and Δ*hisC* pBBR-*oppA2* was observed in the presence of 2 mM of copper by measuring the OD during 48 h (Fig. 5A). Confirming the previous results, Δ*hisC* had a growth defect. The impaired growth due to copper sensitivity was rescued in C6, Δ*hisC*SupPA1 and Δ*hisC* pBBR-*oppA2* strains. These results confirmed the suppressor phenotype in C6 and the role of the mutation identified in this strain to cope with copper stress. Indeed, when reinserted in the Δ*hisC* parent strain, Δ*hisC*SupPA1, the mutation resolved the growth defect observed due to copper sensitivity. Having o*ppA2* on a medium-copy vector allowed bacteria to grow as the WT strain in the presence of copper (Fig. 5A).

**Figure 5.**
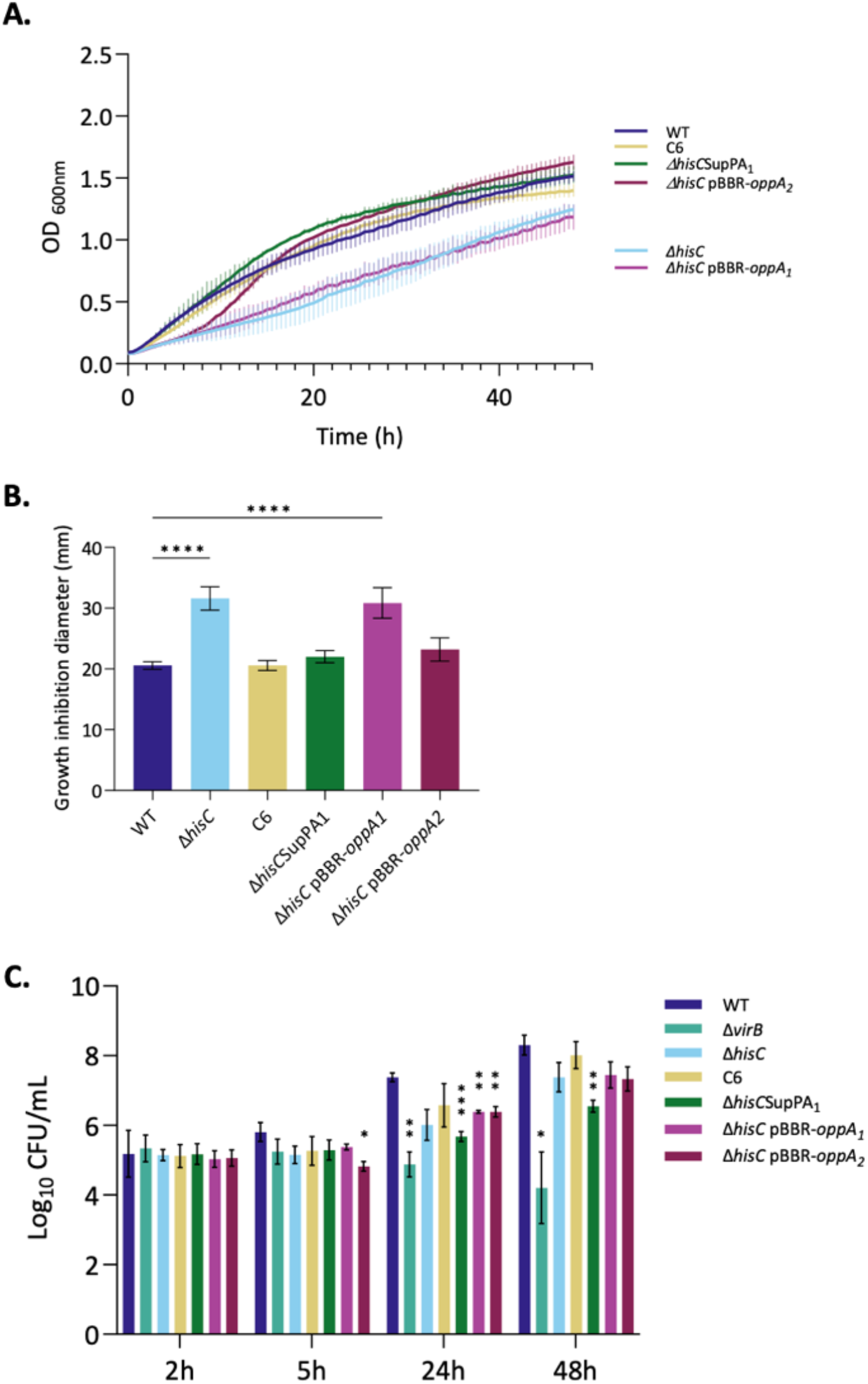
Suppressive mutation saves the sensitivity phenotype to copper stress. WT, Δ*hisC* Δ*hisC*, C6, Δ*hisC*SupPA1, pBBR*-oppA1 and* Δ*hisC* pBBR*-oppA2* strains were tested for their sensitivity to copper toxicity for growth and their intracellular replication capacity. (A) WT, Δ*hisC* strains and derivatives were grown in liquid TSB rich medium containing 2 mM of CuSO_4_. OD _600nm_ was measured every 30 minutes for 48 hours. Data represent three independent replicates. (B) Experiment performed as in Fig. 1C. Data represent the mean ± SD and were compiled from three independent replicates. Statistical analysis was carried out by an ANOVA one-way followed by Dunnett’s multiple comparisons test (****, *P* <0.0001). (C) Intracellular replication of WT, Δ*hisC* mutant and derivatives were assessed by CFU at 2, 5, 24, and 48 hours post-infection of J774.A1 macrophages. Data represent the mean ± SD and were compiled from three independent replicates. Statistical analysis was carried out by an ANOVA two-way followed by Dunnett’s multiple comparisons test (*, *P*<0.05; **, *P* <0.01; ***, *P* <0.001).

The soft agar assay was repeated with these strains and the inhibition zone around the copper well was measured (Fig. 5B). As was evidenced in the case of the liquid growth culture, the C6, Δ*hisC*SupPA1 and Δ*hisC* pBBR-*oppA2* strains exhibited an inhibition zone, akin to that observed in the WT. In contrast, Δ*hisC* and Δ*hisC* pBBR-*oppA1*, both displayed a copper sensitivity.

As an intracellular pathogen, *B. abortus* is subject to various stresses within the cell, including possibly copper intoxication induced by the host cell. We examined the ability of the Δ*hisC* strain and its different derivatives to replicate intracellularly. This was assessed by counting the number of colony-forming units (CFUs) after J774A.1 macrophages infection. The intracellular trafficking of the different strains was monitored at two early time points (2 and 5 hours post infection, PI) and two late time points (24 and 48 hours PI) following replication (Fig. 5C). The WT strain replicated after 24 hours PI (positive control). In contrast, the mutant strain deleted for the entire *virB* operon was used as negative control, since it is known to remain blocked in eBCVs (7,8) and therefore was unable to replicate in macrophages. At 24 hours PI, the *hisC* mutant appeared to have a weaker replication capacity than the WT. However, by 48 hours PI, the CFUs increased and approached the WT level. The C6 suppressor had a replication profile that was slightly higher than that of the *hisC* mutant. Meanwhile, Δ*hisC* pBBR-*oppA1* and Δ*hisC* pBBR-*oppA2* replicated similarly to Δ*hisC*. The strain in which the suppressor mutation was reintroduced, Δ*hisC*SupPA1, appeared to be more affected than the *hisC* mutant itself, particularly at 48 hours. Compared to the *virB* mutant, the *hisC* mutant and its derivatives could replicate within the cell but to a lesser extent than the WT (Fig. 5C).

## Discussion

Copper is an essential transition metal for microorganisms, yet it becomes rapidly toxic at slightly elevated concentration (45,46). In *E. coli*, the total intracellular copper pool (free, labile and sequestered) is around 10^4^ atoms (∼10 μM) where the free cuprous ion amount is at zeptomolar level thus less than one copper atom per cell (15,28). This implies strong buffering systems, likely mediated by cysteine-rich proteins (15,20,45). Metal ions are critical in numerous mechanisms, and their controlled availability is therefore important in the host-pathogen interplay as well. During intracellular pathogenic infection, host cells both deprive pathogens of essential metals such as iron, zinc or manganese, and also exploit copper toxicity as antimicrobial defence, a strategy known as nutritional immunity (13,14,16,17). Notably, IFN-γ-activated macrophages upregulate the copper transporter ATP7A and relocalize it into phagolysosome membrane to poison invading microbes by pumping copper in the pathogen-containing vacuole (13,24). Resistance to copper stress is determinant for virulence in intravacuolar bacteria such as *M. tuberculosis* (25,26) and *S. enterica* (27). Understanding how *Brucella* hold copper homeostasis may be a key to a better understanding to its mechanisms of pathogenicity. In the absence of knowledge regarding the way *Brucella* copes with copper stress, the objective of the present study was to identify potential key actors involved in copper homeostasis through a transposon insertion analysis.

Genes involved in histidine biosynthesis emerged as critical for copper resistance. In this work, the role of this synthesis pathway was therefore investigated through the construction of *hisA, hisB, hisC*, and *hisD* deletion mutants. *Brucella* became auxotrophic for histidine and markedly sensitive to copper, as demonstrated in growth curves and soft agar assays with the loss of one of these genes (Fig. 2). Given the ability of histidine to coordinate metal ions through its imidazole ring, this observation suggests that histidine, free or ligated to a tRNA, might play a role of buffer during copper stress. Our findings therefore revealed an unexpected link between histidine biosynthesis and copper detoxification in *B. abortus*. The copper sensitivity phenotype does not result from a defect in histidine uptake, as supplementation of the minimal medium with histidine restores prototrophy, indicating that the bacterium can import histidine from the extracellular environment (Fig. S4). It has been demonstrated that, amongst a variety of organisms, histidine performs a role in metal resistance or as a buffer. In *Caenorhabditis elegans*, disruption of the histidine ammonia lyase (HALY-1) or dietary histidine supplementation result in an increased of zinc and nickel resistance, possibly through histidine capacity to chelate zinc and nickel (47). In *Aspergillus fumigatus, hisB* deletion decreases resistance to several metals, which can be rescued by histidine supplementation. During iron starvation, an increase in histidine content was observed in this pathogenic fungus (48). Likewise, in *Acinetobacter baumanii*, HutH, the histidine ammonia lyase, increases zinc availability by degrading histidine, which under normal conditions stores zinc in a labile histidine-zinc complex. This finding provides direct evidence for histidine-based buffering mechanisms (49). To support this connection between histidine biosynthesis and metal resistance, a histidine-enrichment analysis of *Brucella* predicted proteome was performed and the results indicated that the majority of proteins exhibiting a histidine percentage exceeding 5% are proteins implicated in metal homeostasis or using metals as cofactors (See Table S1). From this enrichment list, only CueO, the multicopper oxidase, was identified in the Tn-seq analysis performed in the presence of copper, suggesting that histidine-rich proteins cannot buffer copper in a non-redundant manner.

Whole-genome sequencing of the suppressor mutants revealed that, among the suppressors, 15 unique mutations were localized the *opp* operon that encodes an ATP-binding cassette (ABC) transporter, and more specifically the first two genes encoding the periplasmic substrate-binding proteins OppA1 and OppA2. It is striking to observe that no mutations appeared in the permeases *oppB* and *oppC* or the ATPase components, *oppD*/*F*. At least 8 *opp* operons are encoded in *B. abortus* genomes, including 7 on the second chromosome. Among all of them, only one was affected by suppressive mutations within impaired histidine biosynthesis background, highlighting its specificity in rescuing from copper resistance when histidine biosynthesis is lacking. In this study, we demonstrated that when histidine biosynthesis is impaired and the bacteria have to face a copper stress, mutating the two substrate-binding proteins genes, *oppA1* and *oppA2* generates an efficient compensatory mechanism. The mutation in the C6 suppressor is of particular interest due to its location within the promoter region of *oppA*_*1*_. The C6 suppressor and *ΔhisC*SupPA1 strain, in which the mutation was reintroduced into the *ΔhisC* background, were confirmed to display a suppressor phenotype by rescuing growth in the presence of copper in comparison to the parent mutant (Fig. 5A-B). One non-exclusive hypothesis to account for this rescue is that OppA1 and/or OppA2 could facilitate increased histidine uptake, either as free amino acid or within short peptides, thereby enhancing the capacity of the bacterial cytosol to buffer copper ions. In this scenario, the C6 suppressive mutation may lead to *oppA* overexpression. This is consistent with the results obtained with the *oppA1* or *oppA2* overexpression strains, where increasing copy number of the second *oppA2* gene rescued the copper sensitivity (Fig. 5A-B). Increasing the copy number of *oppA1* did not generate the same suppression, maybe because the overexpression of *oppA1* is not achieved in this strain. Nevertheless, the suppressive mutations did not resolve histidine auxotrophy, which makes sense in the absence of histidine or histidine-containing peptides in the medium, if the uptake hypothesis proposed above is true (See Fig. S4). Since the attenuation of *hisC* mutants in infection can be either due to copper sensitivity or histidine auxotrophy, the suppressor strains offer a way to discriminate between these two hypotheses, since the suppressors are copper resistant but still histidine auxotrophic. As indicated in figure 5C, the suppressor mutation in C6 strain does solve the intracellular replication problem of the *hisC* mutant while the suppressive mutation inserted in *hisC* mutant background does not. These results thus demonstrated that within the cell, the replication of *his* mutants is impacted due to a histidine biosynthesis dysfunction rather than a copper stress.

Amino acid starvation is a common feature of host–pathogen interactions. It can be imposed by the host as a nutritional immunity strategy to restrict pathogen growth, or alternatively, it may result from the infection by the pathogen. The source of amino acids for *B. abortus* in host cells is unknown. It may involve extracellular proteases or the hijacking of host proteases to gain small peptides that could fit into the pore of the major porin of *B. abortus*. To the best of our knowledge, such proteases have not been identified so far, and the availability of free amino acids in the rBCV of macrophages is unknown. Such lines of research would deserve an in-depth investigation in the future. Therefore, why is histidine the only amino acid whose biosynthesis would be required in macrophages infection? It is notorious that histidine biosynthesis is a costly amino acid to synthesize in terms of the ATP equivalents need (50). It is therefore possible that the available quantities of histidine in the host cells are a limiting factor for growth. It has been demonstrated that *M. tuberculosis* depends on *de novo* histidine biosynthesis to overcome host-imposed histidine deprivation. IFN-γ-mediated upregulation of histidine catabolizing enzymes (HAL, HDC) reduces free histidine in infected tissues, creating a nutritional stress that *M. tuberculosis* Δ*hisD* mutant cannot withstand. These results highlight the histidine biosynthetic pathway as a critical bacterial strategy in *Mycobacterium* to evade host nutritional immunity and sustain intracellular survival (51). In macrophages, *S*. Typhimurium also rely on histidine biosynthesis genes upregulation to counter free histidine limitation in host cells (52). These studies demonstrate that histidine production is an important adaptation for survival under host-imposed nutrient restriction.

As expected, the results highlighted three genes within the *cue* operon: *copA, cueO* and *cueR* as important for copper resistance *in vitro*. In addition, *copA* transposon mutant was attenuated in macrophages in a previous Tn-seq from our lab (36). Unexpectedly, however, targeted deletions of *copA* or *cueO* did not confer a sensitivity phenotype to copper in our conditions in *B. abortus* (Fig. 1). These results remained consistent when copper concentration was increased. These findings are in striking contrast with observations in *Brucella melitensis* where *bmcO* (*Brucella* multicopper oxidase) mutant displays a copper sensitivity on minimal medium (53), which could be explained by differences related to the bacterial strain context and culture conditions. Also, in *E. coli* and *S*. Typhimurium, the *cue* system is the primary defence line to cope with copper toxicity (28). Indeed, the deletion of *copA* or *cueO* increased copper sensitivity, even stronger under anaerobic conditions for Δ*copA* (31). In *S*. Typhimurium, a similar phenotype has been reported where the loss of *copA* (34) or *cueO* (27) also give a copper susceptibility. Likewise, *cueO* deletion sensitivity phenotype is exacerbated in the absence of oxygen (27). A homologue of *cueO* called *mmcO* also plays a role in copper resistance in *M. tuberculosis* (54). *E. coli* and *S*. Typhimurium have other defence systems, including *cus* and *gol* systems respectively expressed under anaerobic conditions or higher copper concentrations (28,31). Additional copper homeostasis systems are absent in the *Brucella* genome, according to our current knowledge. The role of CopA and CueO was also investigated in *S*. Typhimurium replication capacity within host. While CopA is described having a role in macrophages replication (34), CueO was more important in mouse model (27). This demonstrated the importance of copper homeostasis in *S*. Typhimurium for its virulence. Despite the lack of sensitivity to copper *in vitro*, it would be interesting to test the two mutants constructed in *B. abortus* during macrophage or mice infection to define their role in a more complex virulence model. Even more so as *copA* was identified as required for survival in macrophages(36). It would be also interesting to repeat our experiments in different conditions, such as minimal media. Nevertheless, despite the indications of Tn-seq experiments, the *cu*e system does not appear to be the primary strategy used by *Brucella*, at least under the conditions tested in this work. This could also demonstrate the limitations of Tn-seq analysis in predicting deletion mutant phenotypes.

Our study reveals an intriguing new aspect of copper homeostasis in *B. abortus*. While key copper-associated proteins such as CopA and CueO are critical in other bacteria, they appear surprisingly dispensable in *B. abortus*. Instead, histidine biosynthesis emerges as a possible frontline defence against copper stress. Indeed, we found that disruption of histidine synthesis leads to a copper sensitivity in *B. abortus*, highlighting the importance of this pathway. Interestingly, the attenuation of *hisC* mutant in macrophages stems from histidine auxotrophy rather than copper sensitivity. The mechanism underlying nutrient uptake within the rBCV remain poorly understood, representing a promising avenue for future research into how *B. abortus* acquires amino acids in its intracellular replicative niche.

## Materials and methods

### Bacterial strains and media

*Escherichia coli* DH10B (Invitrogen) andS17-1 (55) strains were grown in Luria-Bertani (LB Lennox) medium at 37°C. *E. coli* MFDpir pXMCS2-Tn*5* (36) was grown in LB medium supplemented with 300 μM *meso-*2,6-diaminopimelic acid. *Brucella abortus* 544 Nal^R^ (referred as the WT in this work; J-M. Verger, INRA, Tours) and its derivative strains were grown in 3% Bacto Tryptic Soy Broth (TSB; Difco™ ref. 211825) rich medium at 37°C. For auxotrophy experiments, a defined medium Plommet Erythritol (PE) (Plommet 1991) was used, composed of 7 or 9.2 g/L K_2_HPO_4_, 3 g/L KH_2_PO_4_, 0.1 g/L Na_2_S_2_O_3_, 5 g/L NaCl, 0.2 mg/L nicotinic acid, 0.2 mg/L thiamine, 0.04 mg/L pantothenic acid, 0.01 g/L MgSO_4_, 0.01 mg/L MnSO_4_, 0.1 mg/L FeSO_4_, 0.1 μg/L biotin and 2 g/L erythritol. All the strains used in this study are list in (Table S5).

When it was necessary, the culture medium was supplemented with the appropriate antibiotics at the following concentrations: Kanamycin (Kan, 10 or 50 μg/ml for chromosomal locus or plasmid selection, respectively), nalidixic acid (Nal, 25 μg/ml).

### Strains construction

For deletion mutants, the whole gene was removed by homologous recombination. Joined PCR was applied. Two regions (the upstream and downstream regions of the target gene) of more than 500 base pairs were amplified from purified *B*. abortus 544 gDNA using Q5® High-Fidelity DNA Polymerase (New England Biolabs) and primer pairs (F1/R1 for the upstream region and F2/R2 for the downstream region). The two amplified fragments were fused together through a complementary region designed in the primers used and the resulting fragment was amplified by PCR using F1 and R2 primers. To Δ*hisC*SupPA1 construction, the complementary region primer pair contained the desired SNP. The amplicon was purified and inserted in an *Eco*RV (New England Biolabs)-linearized pNPTS138 plasmid through an overnight ligation (T4 DNA ligase, Promega) at 20°C. The ligation product was transformed in DH10B *E. coli* and clones positive in blue-white screening were screened by PCR using GoTaQ® DNA polymerase (Promega). Selected plasmid was purified and checked by sequencing. The plasmid was inserted in S17-1 *E. coli* to allow conjugation to *B. abortus* 544 Nal^R^ by mating. Allelic exchange on the chromosome occurs *via* the non-replicative plasmid, pNTPS138 (56). The gene deletion was then verified by PCR using GoTaQ® DNA polymerase (Promega). Δ*hisA*, Δ*hisB*, Δ*hisC* and Δ*hisD* deletion strains were complemented with pMR10 carrying the deleted gene. Complementation plasmids were obtained by amplification of the genes with a region of 400 base pairs upstream the coding sequence by PCR using Q5® High-Fidelity DNA Polymerase (New England Biolabs). The PCR product was digested as well as the pMR10 plasmid with BamHI and XbaI enzyme (New England Biolabs). The insert was inserted in the linearized plasmid through ligation in the same direction as the *E. coli lac* promoter of the vector.

To construct the overexpression strains, the genes were amplified by regular PCR using Q5® High-Fidelity DNA Polymerase (New England Biolabs). The designed primers included restriction sites for KpnI and SacI restriction enzymes (New England Biolabs). PCR product was purified and restricted by KpnI and SacI as well as the multicopy-plasmid pBBR-MCS2 (56). The gene expression is under the control of the *E. coli lac* promoter.

All the primers and plasmids used in this study are listed in Table S3 and S4 respectively.

### Transposon sequencing assay

One millilitre of an overnight culture of *B. abortus* 544 Nal^R^ and 50 μl of overnight culture of *E. coli* MFDpir pXMCS2-Tn*5* Kan^R^ were mixed. After recovery in 1mL of TSB supplemented with *meso-*2,6-diaminopimelic acid (300 μM), OD of the cultures was measured and normalized to a DO of 1. Using a 96-well plate, serial 10-fold dilutions were carried out. Fifteen μL of these dilutions were plated on TSB plates supplemented with CuSO4 concentrations ranging from 0.5mM to 3mM. CFUs were counted to determine the optimal concentration of copper. A CuSO_4_ concentration of 2mM was selected. The mating was repeated as described above, and plates were incubated overnight at RT. To ensure complete coverage of the genome, 8 *B. abortus* and *E. coli* matings were done in parallel. The matings were recovered in 1mL of TSB from which 50 μl were diluted in 450 μl of TSB. One hundred μl of the dilution were spread on TSB agar plates supplemented with kanamycin or kanamycin and CuSO_4_ to allow the selection the resulting *B. abortus* mini-Tn*5* libraries. After 4 days of incubation, the *B. abortus* mutants were recovered in 2 ml of TSB. The bacterial suspension was centrifugated for 7 minutes at 7000 RPM. The pellet was resuspended in 300 μl of 2% SDS. Bacteria were inactivated at least 1 hour at 80°C. All the samples were pooled together following the two conditions. gDNA was extracted using the Nucleobond AXG 500 kit (Macherey-Nagel). This gDNA was sequenced via Illumina sequencing. The raw data were analysed by an automatic process TnBox (https://github.com/fxstubbe/TnBox), following a previously described method (37). TnBox uses BWA method (57) to map the raw reads from Illumina sequencing on the genome of *B. abortus* 544 and the SAMtools suite (58) to calculate read counts. The libraries underwent normalization based on read-depth. For a detailed protocol see ref (37). From the library, 4.8 x 10^6^ random mutants were recovered and sequenced using a deep sequencing method. We identified 1 684 839 and 1 743 143 unique insertion sites for control and copper condition respectively, illustrating genome saturation with a unique insertion every 1.95 bp for control and every 1.89 bp for the stress condition, on average. To evaluate the contribution of each gene to *Brucella* fitness under copper stress, a transposon insertion frequency (TnIF) was calculated. The TnIF was defined as the average log_10_(r+1)/l, where *r* is the number of miniTn*5* insertions at a given nucleotide and *l* is the coding sequence length in base pairs. To minimize insertional bias, the calculation was performed considering only the central 80% of the coding sequence. For each gene, the differential TnIF (ΔTnIF) between interest and control condition was calculated as ΔTnIF = TnIF*copper* – TnIF*control*. A negative ΔTnIF indicates that the gene is required in the copper condition as the number of reads associated is lower than the control condition.

### Growth curves

Growth was assessed by using BioTek Epoch2 Microplate Reader by measuring OD_600nm_ every 30 minutes for 48 hours at 37°C with agitation. Culture of *B. abortus* 544 and derivative strains were grown in TSB overnight to reach exponential phase (OD_600nm_ 0.3-0.8) and normalised at OD_600nm_ 0.1 final the following day. For the growth measurement in presence of copper, TSB was supplemented with 1.6 or 2 mM of CuSO_4_. For the growth measurements in PE, overnight bacterial cultures were wash twice in PBS and diluted to an OD_600nm_ of 0.1 in PE. When necessary, PE was supplemented with 1 mM of histidine (Sigma).

### Suppressor assays

Culture of *B. abortus* 544 and derivative strains were grown in TSB overnight to reach exponential phase (OD_600nm_ 0.3-0.8) and normalised at OD_600nm_ 0.1 . Twenty μl of each strain were spread on TSB agar plate containing 1.6 or 2 mM CuSO_4_. Plates were incubated at 37°C for 4 to 5 days.

For all four Δ*his* mutants, 29 isolated colonies formed on CuSO4 plates were streaked on new TSB agar 1.6 and 2 mM CuSO4 plates to confirm the phenotype of the suppressors. Among the 29 clones, 21 were still resistant to copper on the new plates and were grown overnight and stored. To extract demonic DNA, bacterial cultures of around 7.5 ml were centrifuged at 8200 x *g* for 5 minutes. Supernatants were discarded, and bacterial pellets were resuspended in 300 μl of PBS. Then, bacteria were inactivated at 80°C during at least 1 hour. Afterwards, 100 μl of SDS 10 % were added. The gDNA extraction was achieved following manufactural instructions of the Macherey-Nagel™ NucleoSpin™ Tissue Kit. Bacterial SNP sequencing was performed by BIO, part of Pathology and Genetics Institute (http://www.bio-be.be/biopharma-croservices/). The analysis of sequencing reads was conducted utilising the Snippy tool, which is available on the Galaxy.org website (https://usegalaxy.org/). Each parental mutant and suppressor genome was aligned to the reference *B. abortus* 544 genome.

### Soft agar well diffusion assays

Culture of *B. abortus* 544 and derivative strains were grown in TSB overnight to reach exponential phase (OD_600nm_ 0.3-0.8) and normalised at OD_600nm_ 0.1 Five hundred μl were added to 4.5 ml of TSB soft agar (0.7% agar) and spread on a 20 ml TSB agar plate. After the culture-soft agar mix had dried, a well was dug the upper part of a pipet tip and removed. The well was filled with 100 μlof 200 mM CuSO_4_. The plates were incubated for 3-4 days at 37°C upward-facing lid the first day before being flipped upside down. After incubation, the inhibition zone around the wall was measured in three different directions and the mean was reported in a graph in millimetres.

### J774A.1 macrophages culture and infection

J774A.1 macrophages (ATCC) were cultivated at 37°C in a 5 % CO2 atmosphere in GlutaMAX™-supplemented Dulbecco’s Modified Eagle Medium (DMEM, Gibco™) in which 10 % of heat inactivated fetal bovine serum (Gibco™) was added. The day before infection, J774A.1 macrophages were seeded in a 24-wells plate at a concentration of 1.10^5^ cells/ml. On the day of the infection, overnight cultures of *Brucella* strains at exponential phase (0.3-0.8) were washed twice in PBS and diluted in GlutaMAX™-supplemented DMEM medium at a multiplicity of infection (MOI) of 50. Bacteria suspensions were added to the cells, and the 24-wells plates were centrifuged at 169 x *g* for 10 minutes at RT. Infected cells were incubated at 37°C in a 5 % CO2 atmosphere for 1 hour. Cell medium was refreshed and supplemented with 50 μg/ml gentamycin to kill extracellular bacteria. Cells were incubated one extra hour before media was again refresh and supplemented with 10 μg/ml gentamycin, to kill bacteria egressed and avoid re-infection event, for the 46 hours left. For CFU counts, at interested time point (2,5, 24 or 48 hours after infection) cells were washed twice with PBS and incubated with 0.1% Triton X-100 PBS for 10 minutes at RT before being scratched to be detached. Macrophage lysates were 10-fold diluted in PBS and specific dilution (20 μl) were spread onto TSB agar plates. Plates were incubated at 37 °C for 5 days and CFU were counted. The CFU number (log10) was calculated per ml of lysate and reported into a graph.

### Statistical analysis

Statistical analyses *via* one-, or two-way ANOVA with Dunnett’s test were carried out provided by the GraphPad Prism software. Values of p < 0.05 were considered to represent a significant difference.

## Acknowledgements

The authors thank the technical team of the URBM for their support, as well as the University of Namur for the access to the BL3 platform, and the logistic and financial supports. The authors thank Damien Devos (Pasteur Lille) for the analysis of histidine content in the predicted proteome of *B. abortus*, and François Jacob-Dubuisson (Pasteur Lille) and Hilde De Reuse (Pasteur Paris) for stimulating discussions.

C.F. was supported by a FRIA (FNRS) PhD fellowship. A.R. was supported by a Aspirant (FNRS) PhD fellowship. K.P. was supported by a Chargé de Recherche grant fellowship from FNRS. This publication is supported by the Walloon Region as part of the funding for the FRFS-WELBIO strategic axis (X.1512.24). The work was also supported by PDR grants T.0058.20 and T.0068.24 from FRS-FNRS, as well as Concerted Research Action 17/22-087 and 22/27-128 from the Fédération Wallonie-Bruxelles.

